# Two distinct modes of meiotic chromosome synapsis

**DOI:** 10.64898/2026.08.19.745786

**Authors:** Lauren M. Lotka, Amy J. MacQueen, Carolyn R. Milano, Nancy M. Hollingsworth, Andreas Hochwagen

## Abstract

The pairwise alignment of homologous chromosomes within the synaptonemal complex (SC) is important for meiotic crossover recombination and fertility ^1–3^. However, chromosomes do not need sequence homology to synapse, with meiotic recombination defects often leading to synapsis of non-homologous chromosome segments ^2,4,5^. Here we show that such heterologous synapsis reflects a distinctly regulated mode of meiotic chromosome synapsis that also happens during the early stages of wild-type yeast meiosis and occurs in parallel to the well-known synapsis initiation at crossover-designated sites ^3,5,6^. Heterologous synapsis initiates along chromosome arms after double-strand break resection and is accompanied by canonical markers of crossover repair, but does not need recombinase-dependent strand invasion. Instead, it requires the DNA-damage sensor kinase ATR/Mec1, which phosphorylates of a specific amino acid in Zip1, the major transverse filament protein of the SC. Phospho-mimetic mutants in *ZIP1* rescue the synapsis defect of *mec1* mutants and also partially restore gamete viability, indicating that this particular *MEC1* function is important for the faithful completion of meiosis. Importantly, crossover repair quickly rectifies heterologous synapsis, allowing successful completion of meiosis even when most of the genome initially synapses independently of homology. Our data identify meiotic chromosome synapsis as a dynamic and reversible process that becomes optimized as result of recombination-dependent chromosome pairing.

---

Meiosis is a specialized cell division that produces haploid gametes from diploid germ cells. To reduce the genetic material, all chromosomes must identify their homologous partners. In many organisms, including mammals, this process requires the formation of programmed DNA double-strand breaks (DSBs) and their repair into crossovers. Meiotic pairing typically coincides with the formation of the proteinaceous synaptonemal complex (SC) on chromosomes ^1–3^. The SC is a highly conserved tripartite structure consisting of lateral elements, formed by the axes of aligned chromosomes, and a central region, comprised of a central element and transverse filaments that bridge the lateral and central elements (**Fig. 1a**). In the yeast *Saccharomyces cerevisiae*, the transverse filaments consist of homodimers of the coiled-coil protein Zip1 that are oriented perpendicularly to the axis and connect to Gmc2 and Ecm11 in the central element ^7,8^. The SC has multiple functions, including promoting meiotic crossover recombination, preventing excess DSBs, and the temporal control of repair patterns ^3,5,9^. Failure to form SCs can lead to the mis-segregation of homologous chromosomes and gametic aneuploidy ^10,11^.

**Figure 1:**
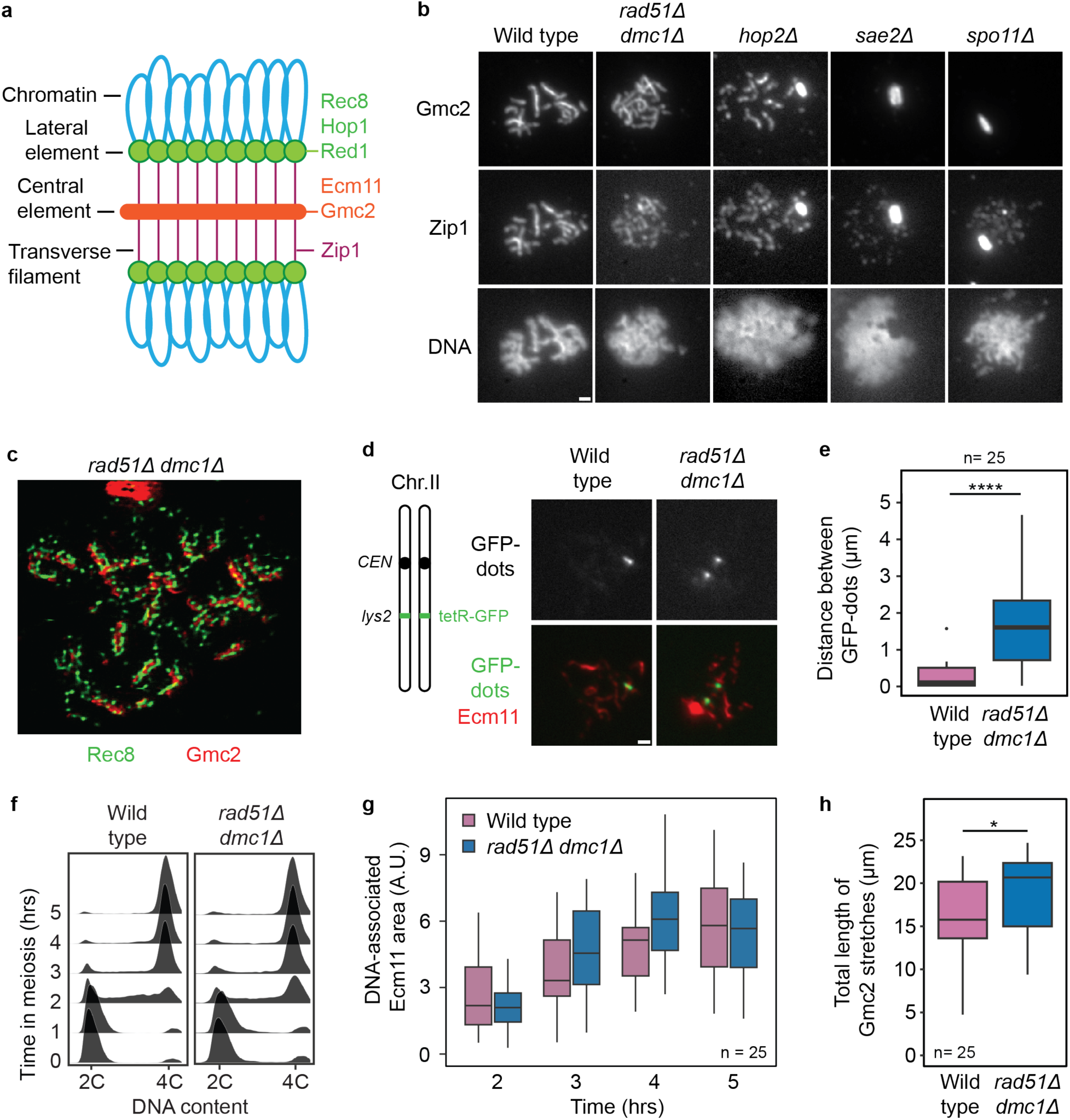
Chromosome synapsis occurs in the absence of strand invasion. **a**, Schematic of the SC, highlighting major structural elements, as well as key protein components in *S. cerevisiae*. **b**, Spread nuclei from synchronous cultures of wild type, *rad51Δ dmc1Δ*, *hop2Δ*, *sae2Δ*, and *spo11Δ.* Immunostaining was performed with antibodies for Gmc2 and Zip1 at hour 4. DNA was counterstained with DAPI. **c**, Immunostaining of a surface-spread *rad51Δ dmc1Δ* nucleus at hour 4 of a synchronous meiotic culture. Chromosomes were stained using antibodies for Rec8 (lateral element) and Gmc2 (central element). Imaging was done with STED microscopy. **d,** (left) Schematic of the GFP-dots pairing assay. (right) Immunostaining was performed on spread nuclei at hour 4 with an antibody for Ecm11 (central element). **e**, Distance between GFP-dots as a measure of pairing in wild type and *rad51Δ dmc1Δ* at hour 4. Box-and-whisker plots show medians and interquartile ranges. **f**-**g**, Analysis of progressive Ecm11 installation in a synchronous meiotic time course of wild type and *rad51Δ dmc1Δ*. **f**, Synchrony of prophase entry as determined by DNA content analysis, which measures bulk DNA replication in premeiotic S phase. **g**, Quantification of DNA-associated Ecm11 area (arbitrary units, A.U.) per nucleus in wild type at hours 2 (2.53 ± 1.57), 3 (3.81 ± 2.16), 4 (4.96 ± 1.70), 5 (6.07 ± 2.52) and *rad51Δ dmc1Δ* at hours 2 (2.11 ± 1.01), 3 (4.63 ± 2.14), 4 (6.22 ± 2.04), 5 (5.26 ± 2.24). The differences at each time point are not significant. **h**, Quantification of the total length of Gmc2 stretches (μm) in wild type (16.20 ± 4.38) and *rad51Δ dmc1Δ* (19.10 ± 5.50) at hour 4. All experiments were repeated at least twice with similar results. The data are presented as mean ± standard deviation (s.d.). Scale bar, 1 μm. Significance was evaluated using Mann-Whitney U test. *P ≤ 0.05, ****P ≤ 0.0001

SCs typically connect homologous chromosome pairs because SC assembly is coupled to crossover-fated DNA repair ^3,5^. However, when crossover repair is impeded, SC structural components can form linear assemblies between non-homologous chromosome axes ^2,4,5^. SC can even form between sister chromatids ^12^. We refer to situations where SC components assemble into linear structures independently of the underlying sequence homology as heterologous synapsis. In *S. cerevisiae*, such heterologous synapsis is readily observed in haploid cells undergoing meiosis ^13^ and in mutants lacking the recombinase proteins Dmc1 and/or Rad51 ^14–16^ or the recombinase co-factors Hop2 or Mnd1 ^17,18^. However, the mechanisms triggering heterologous synapsis are unknown.

It is also unclear whether heterologous synapsis is a terminal pathology or whether it represents a normal intermediate of meiotic chromosome synapsis that is ultimately corrected. The latter notion is supported by work showing that the SC dynamically assembles and disassembles and shares features with liquid-liquid phase-separated systems ^5,19,20^. In addition, electron microscopy analyses of chromosomally abnormal karyotypes indicate that the SC has the capacity for partner switches, as initially correct synapsis can become non-homologous ^2,21^. Moreover, work in hexaploid wheat and other plants suggests that the converse may also be true, namely that initially incorrect synapsis can be corrected ^2^.

To study the regulation of heterologous synapsis, we initially focused on situations where recombinase-dependent chromosome pairing is prevented. *S. cerevisiae hop2Δ* mutants display extensive heterologous synapsis ^17^ but because both recombinases remain present, a background level of homologous recombination cannot be excluded. Therefore, we also analyzed *rad51Δ dmc1Δ* mutants, which lack both recombinases and were recently shown to accumulate the transverse filament protein Zip1 on chromosomes ^16^ (**Fig. 1b**). Immunostaining of chromosome spreads of meiotic *rad51Δ dmc1Δ* cells revealed the expected tripartite SC structure, comprised of two lateral elements, containing the axis protein Rec8, and a central element, marked by Gmc2 (**Fig. 1c**) ^7,8^. Moreover, Zip1 localized along linear assemblies of the central element protein Ecm11 (**Fig. 1b**; **Extended Data, Fig. 1a**). Central-element assembly depended on *ZIP1* and did not require the strand-annealing factor *RAD52*, excluding Rad52-dependent homologous recombination as a trigger for synapsis initiation (**Extended Data, Fig. 1b**). Analysis of chromosomally encoded GFP marks (GFP dots) confirmed that the linear assemblies of SC proteins frequently formed along chromosomes that are not engaged with their homologous partner (**Fig. 1d-e**). Notably, central element tracks (Ecm11-Gmc2) formed with wild-type-like kinetics in *rad51Δ dmc1Δ* mutants (**Fig. 1f-g**) and the maximal lengths of central-element tracks as well as total occupied chromosomal area were comparable to wild type, both in normally progressing and in prophase-arrested cells (*ndt80Δ*; **Fig. 1h**; **Extended Data, Fig. 1c-d**). Thus, *rad51Δ dmc1Δ* mutants undergo extensive heterologous synapsis in the absence of strand invasion.

We sought to determine which signals trigger heterologous synapsis. Although certain strain backgrounds allow synapsis without meiotic DSBs ^22,23^, the synapsis observed in *rad51Δ dmc1Δ* mutants depended fully on the DSB-inducing enzyme Spo11 (**Extended Data, Fig. 1b**). Central element signal was also absent in *sae2/com1Δ* mutants (**Fig. 1b**), which fail to remove Spo11 from break ends and do not initiate break resection ^24–26^, indicating that synapsis requires free DNA ends and/or exposed single-stranded DNA (ssDNA). Centromeres can serve as sites of SC formation ^27^. However, wild-type and *rad51Δ dmc1Δ* chromosomes displayed a similar number of linear central element assemblies without associated centromere signals (**Fig. 2a-b)**, indicating that heterologous synapsis can nucleate on chromosome arms. Notably, analysis of chromosome spreads showed approximately 16 kinetochore foci in *rad51Δ dmc1Δ* cells, rather than the 32 expected if centromeres/kinetochores were unpaired (**Extended Data, Fig. 2a-b**). This number likely reflects the coupling of non-homologous centromeres that occurs independently of DSB formation ^28^.

**Figure 2:**
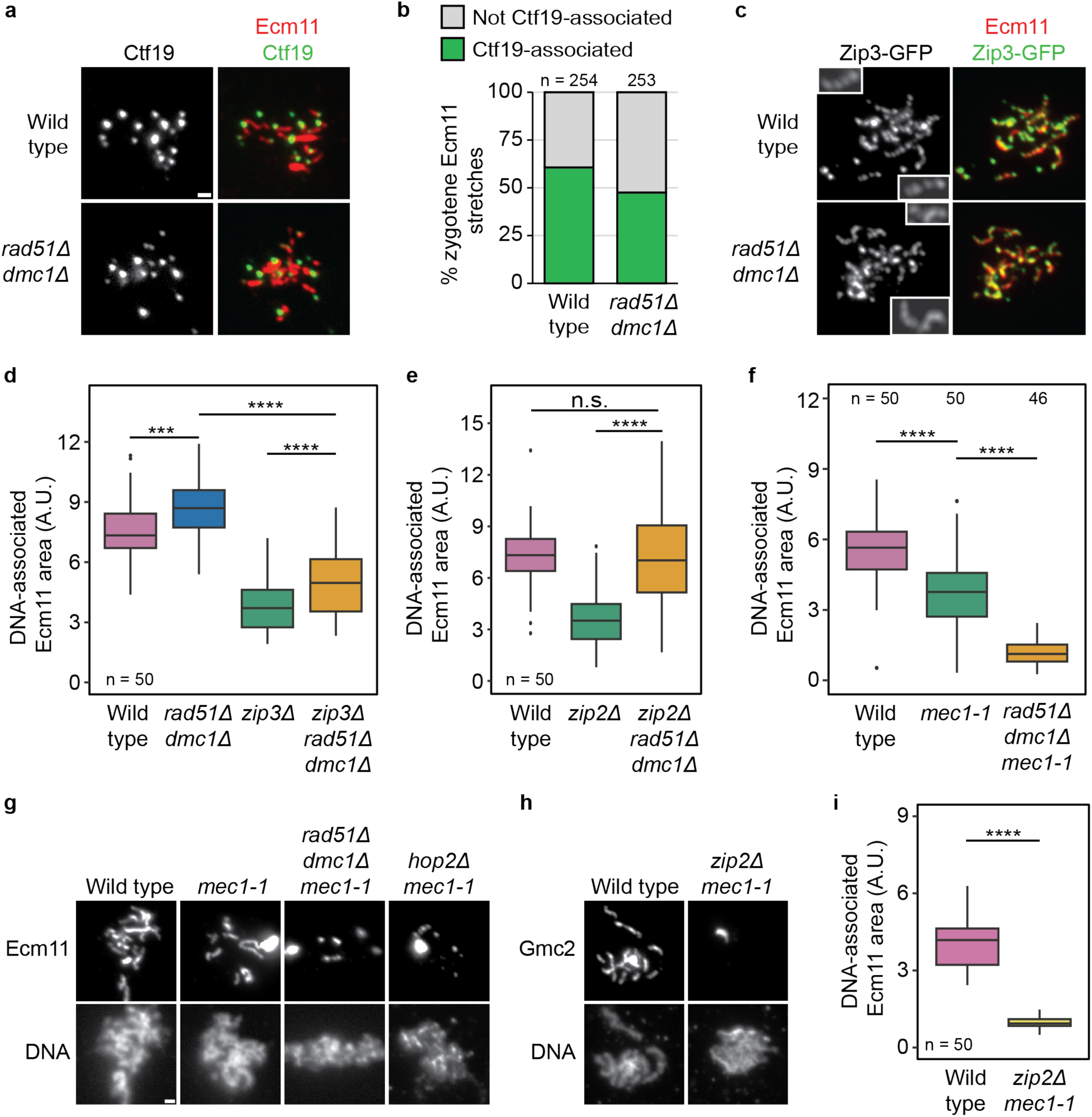
Heterologous synapsis depends on *MEC1*. **a**, Immunostaining of spread wild-type and *rad51Δ dmc1Δ* nuclei was performed at hour 3 of synchronous meiotic time course with antibodies for Ecm11 and the kinetochore protein Ctf19. **b**, Ecm11 stretches on zygotene nuclei (characterized by short SC stretches) that overlapped with Ctf19 foci were counted as centromere-associated (WT: 154/254, *rad51Δ dmc1Δ*: 120/253), Ecm11 stretches with no overlapping Ctf19 foci were counted as not centromere-associated. **c**, Analysis of Zip3-GFP foci on spread chromosomes from hour 4. Immunostaining was performed with antibodies for Ecm11 and GFP. Insets highlight examples of Zip3 foci that are qualitatively different between WT and *rad51Δ dmc1Δ*. **d**, Quantification of DNA-associated Ecm11 area (A.U.) at hour 4 in wild type (7.63 ± 1.55), *rad51Δ dmc1Δ* (8.81 ± 1.73), *zip3Δ* (3.88 ± 1.55), and *zip3Δ rad51Δ dmc1Δ* (4.96 ± 1.62); see **Extended Data, Fig. 2f** for representative images. **e**, Quantification of DNA-associated Ecm11 area (A.U.) at hour 4 in wild type (7.35 ± 1.80), *zip2Δ* (3.59 ± 1.70), and *zip2Δ rad51Δ dmc1Δ* (7.31 ± 3.40); see **Extended Data, Fig. 2g** for representative images. **f**, Quantification of DNA-associated Ecm11 area (A.U.) in wild type (5.59 ± 1.39), *mec1-1* (3.69 ± 1.63), and *mec1-1 rad51Δ dmc1Δ* (1.19 ± 0.53). Immunostaining was performed at hour 4 with an antibody for Ecm11. Note: all *mec1-1* strains in this study were also *sml1Δ* to support viability of *mec1-1* mutants. **g**, Representative images of data in (f). DNA was counterstained with DAPI. **h**, Immunostaining of wild type and *zip2Δ mec1-1* was performed at hour 4 with an antibody for Gmc2. DNA was counterstained with DAPI. **i**, Quantification of DNA-associated Ecm11 area (A.U.) in wild type (3.97 ± 0.99) and *zip2Δ mec1-1* (1.38 ± 1.96). The data presented are from chromosome spreads of synchronous meiotic yeast cultures and presented as mean ± s.d. Scale bars, 1 μm. Significance was evaluated using Mann-Whitney U test. n.s., not significant. ***P ≤ 0.001, ****P ≤ 0.0001.

Consistent with synapsis initiation on chromosome arms, we observed abundant foci of the synapsis regulator Zip3 on *rad51Δ dmc1Δ* chromosomes (**Fig. 2c**). Zip3 foci are also observed in *dmc1Δ* and *rad51Δ* single mutants ^29,30^. While total Zip3 signal on *rad51Δ dmc1Δ* chromosomes was similar to wild type, focus number was reduced, with Zip3 foci often appearing larger and more elongated in *rad51Δ dmc1Δ* (**Fig. 2c**, insets; **Extended Data, Fig. 2c-d**). The crossover repair factor Msh4 also formed foci on *rad51Δ dmc1Δ* chromosomes (**Extended Data, Fig. 2e**), indicating that at least two markers of crossover designation and repair become locally enriched on chromosomes independently of strand invasion.

Deletion of *ZIP3* reduced synapsis similarly in *rad51Δ dmc1Δ* and wild-type backgrounds (**Fig. 2d**; **Extended Data, Fig. 2f**), suggesting that despite their abnormal morphology, Zip3 foci in *rad51Δ dmc1Δ* mutants are competent for synapsis initiation. On the other hand, heterologous synapsis did not require the crossover-associated synapsis initiation factor Zip2 ^31^ (**Fig. 2e**; **Extended Data, Fig. 2g**). These data are consistent with the fact that meiotic crossover repair does not occur in *rad51Δ dmc1Δ* mutants ^32^ and reveal different activities of these two synapsis initiation factors, possibly linked to the fact that Zip2 and Zip3 interface with different SC components ^6,33^. These results indicate that initiation of heterologous synapsis is genetically separable from homologous synapsis.

In keeping with heterologous synapsis being distinctly regulated, several structural features distinguished it from wild-type SC. For instance, although the fluorescence intensity and area of the Ecm11 signal of *rad51Δ dmc1Δ* chromosomes were comparable to wild type, the fluorescence intensity was only about half of wildtype and signal area of Zip1 was reduced (**Extended Data, Fig. 1c, 3a**). As central element components are continuously deposited on chromosomes ^8,34^, these data suggest that Zip1 deposition is uncoupled from central element assembly when synapsis occurs independently of homology search. In line with heterologous SC not maturing like wild-type SCs, chromosomal association of the axis protein Hop1, a hallmark of chromosomes in early prophase ^35^, remained high and more continuous on synapsed chromosomes in *rad51Δ dmc1Δ* mutants (**Extended Data, Fig. 3b-c**). Thus, heterologous synapsis is structurally different from mature SC.

We aimed to identify factors that specifically regulate heterologous synapsis. The requirement for resected DSBs (**Fig. 1b**) prompted us to investigate a role of Mec1^ATR^ kinase, which is activated by ssDNA ^36–38^. Immunostaining showed that while synapsis is partly defective in a *mec1-1* mutant, which truncates the Mec1 kinase domain ^39,40^, central element assembly is almost completely abolished in *rad51Δ dmc1Δ mec1-1* mutants (**Fig. 2f-g**). These data imply the existence of two synapsis initiation pathways, one activated by Mec1 in response to resected DSBs, the other activated by Zip2 and its interactors in response to crossover repair. Consistent with this interpretation, *mec1-1* and *zip2Δ* single mutants displayed partial synapsis defects in otherwise wild-type cells (i.e. *RAD51 DMC1*; **Fig. 2e-f, Extended Data, Fig. 2g**), whereas synapsis was minimal in *mec1-1 zip2Δ* double mutants (**Fig. 2h-i**). We conclude that *S. cerevisiae* employs two pathways to initiate synapsis in response to meiotic DSBs.

One of the targets of Mec1 kinase activity is Zip1 serine 75 (S75) ^41^. This phosphorylation mark primes Zip1 for further phosphorylation and is important for the regulation of Zip1-dependent centromere coupling ^41^. When S75 was replaced with alanine (*zip1-S75A*) in otherwise wild-type cells, synapsis proceeded with no obvious defects (**Fig. 3a**) ^41^. However, S75 was essential for synapsis in the *rad51Δ dmc1Δ* background. These data indicate that Mec1 acts through Zip1-S75 to control heterologous synapsis.

**Figure 3:**
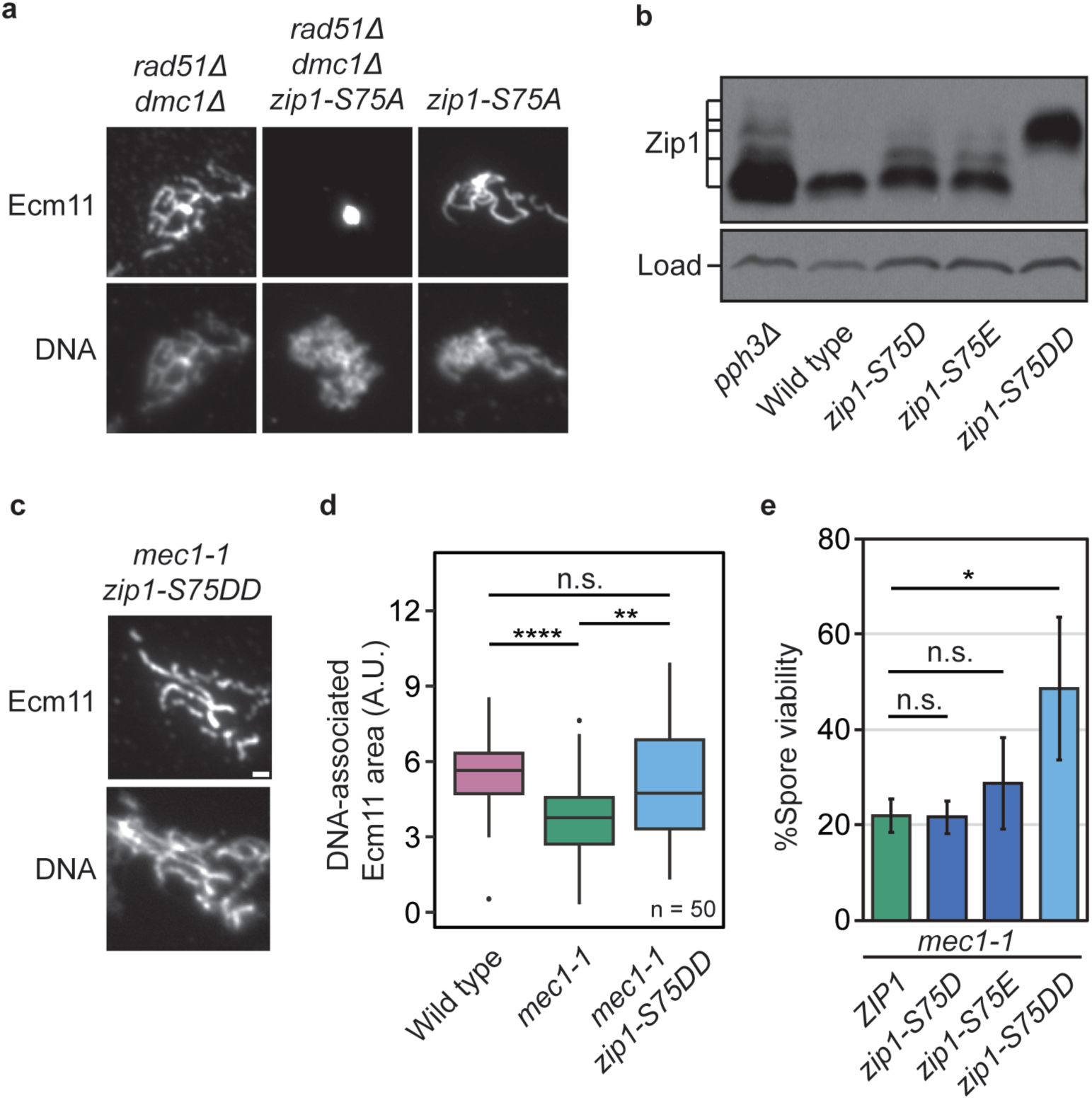
MEC1-dependent synapsis requires phosphorylation of Zip1-S75. **a**, Immunostaining of spread nuclei of *rad51Δ dmc1Δ*, *rad51Δ dmc1Δ zip1-S75A*, and *zip1-S75A* was performed at hour 4 of synchronous meiotic time course with an antibody for Ecm11. DNA was counterstained with DAPI. **b**, Western blot samples collected at hour 3 were probed for Zip1. All higher migrating species depend on Zip1-S75 because Zip1-S75 phosphorylation acts as a priming event for further phosphorylation ^41^. A cross-reacting band served as loading control (load). Zip1-S75DD denotes Zip1-L74D, S75D. **c**, Immunostaining of spread *mec1-1 zip1-S75DD* nuclei was performed at hour 4 with an antibody for Ecm11. DNA was counterstained with DAPI. **d**, Quantification of DNA-associated Ecm11 area (A.U.) at hour 4 in wild type (5.59 ± 1.39), *mec1-1* (3.69 ± 1.63), and *mec1-1 zip1-S75DD* (5.06 ± 2.21). The data and presented as mean ± s.d. Scale bars, 1 μm. Significance was evaluated using Mann-Whitney U test. n.s., not significant, **P ≤ 0.01, ****P ≤ 0.0001. **e**, Quantification of percent spore viability in *mec1-1 ZIP1* (22 ± 3), *mec1-1 zip1-S75D* (22 ± 3), *mec1-1 zip1-S75E* (29 ± 10), and *mec1-1 zip1-S75DD* (49 ± 15) strains. Significance was evaluated using a two-tailed Welch’s t-test. n.s., not significant, *P ≤ 0.05.

To test whether phosphorylation of Zip1-S75 is also sufficient to initiate *MEC1*-dependent synapsis, we constructed a series of phospho-mimetic mutations by replacing S75 with either aspartate (D) or glutamate (E). To better mimic the two negative charges of the phosphate group, we also introduced two adjacent aspartates (L74D,S75D) ^42^. Western blot analysis showed that the S75D and S75E mutations were partially effective in priming Zip1 phosphorylation, as evident from a ladder of slower-migrating species that is also seen when Zip1 dephosphorylation is suppressed by eliminating the Pph3 subunit of PP4 phosphatase (**Fig. 3b**) ^41^. By contrast, the L74D,S75D mutation (*zip1-S75DD*) led to a quantitative shift to the highest migrating species, indicating highly efficient priming. Importantly, the *zip1-S75DD* mutation was sufficient to rescue the synapsis defect (**Fig. 3c-d**) and partially restore spore viability of a *mec1-1* mutant (**Fig. 3e**), whereas the *zip1-S75D* and *zip1-S75E* alleles had no significant effects on spore viability, mirroring the weaker phosphorylation priming (**Fig. 3b**). These data indicate that Zip1-S75 phosphorylation is a key target for Mec1-dependent synapsis initiation and is important for gamete survival.

Although resected DNA ends accumulate in *rad51Δ dmc1Δ* and *hop2Δ* mutants ^32,43^, such ends are also normal intermediates of meiotic recombination, implying that heterologous synapsis may also occur in wild-type cells. Indeed, 64% of unpaired chromosomal GFP dots in wild-type zygotene nuclei overlapped SC signal, as defined by short linear assemblies of central element proteins (**Fig. 4a-b**). This number is significantly higher than random (p=0.0019, t-test). Moreover, the fraction of unpaired GFP-dots overlapping central element structures in zygotene was significantly less in *mec1-1* mutants (**Fig. 4b**; p=0.0022, t-test), as expected from *MEC1*-dependent activation. These data suggest that SC structures can transiently join heterologous chromosomes during wild-type meiosis. As wild-type chromosomes ultimately synapse with their homologous partners, these data also imply that heterologous synapsis is reversible, perhaps via transient disassembly as previously observed in *zip3* mutants ^19^.

**Figure 4:**
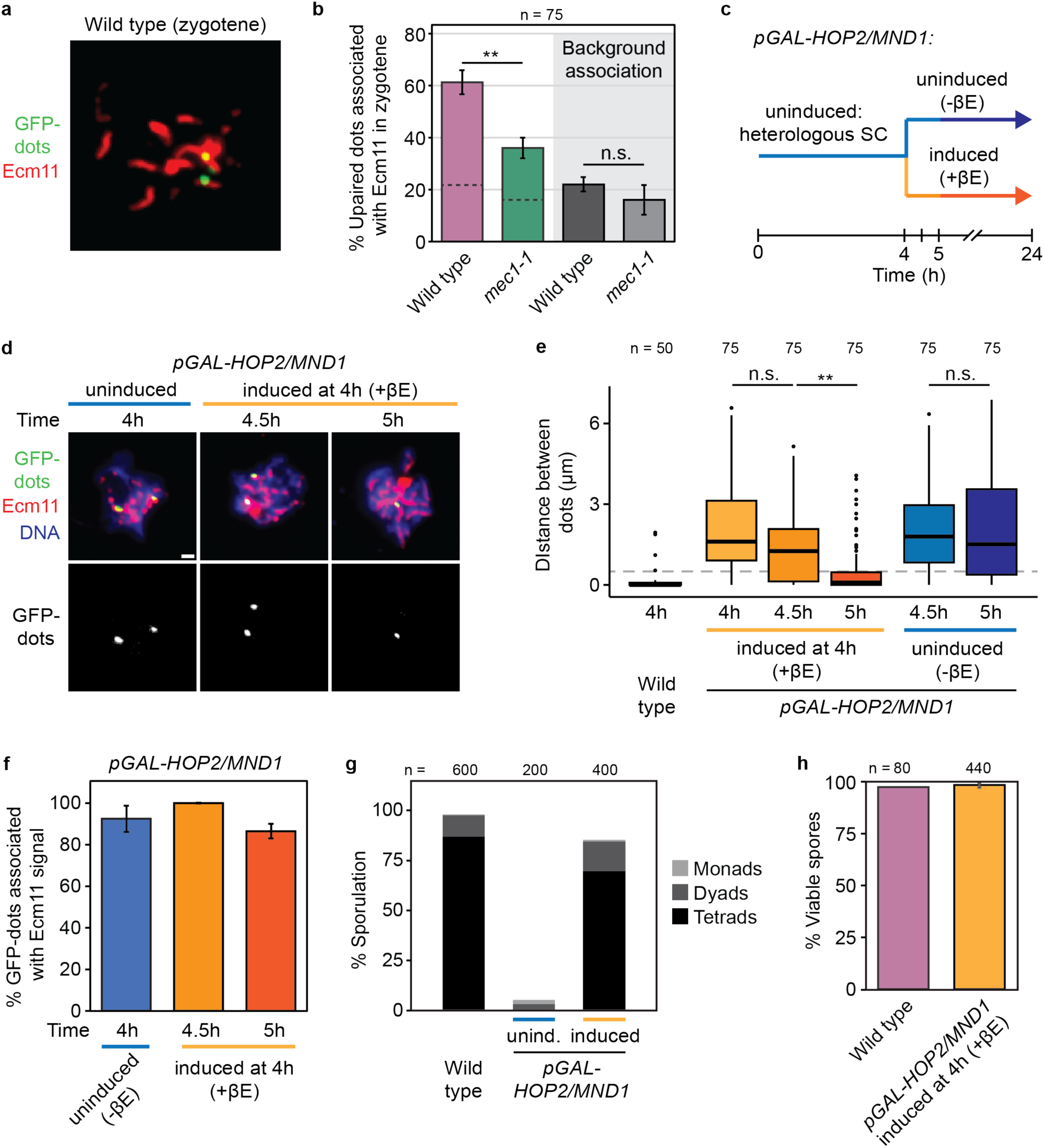
Repair-mediated correction of heterologous synapsis supports gametogenesis. **a**, Immunostaining of spread wild-type nuclei was performed at hour 3 of a synchronous meiotic time course with antibodies for Ecm11 and GFP. **b**, Quantification of percent of unpaired dots associated with Ecm11 in zygotene at hours 2 and 3 in wild type (61.30 ± 4.62) and *mec1-1* (36 ± 4) and in background association control where the Ecm11 image was rotated 90 degrees with respect to the GFP image in wild type (22 ± 2.83) and *mec1-1* (16.00 ± 5.66). Pairing was defined as a distance of dot centers of ≤ 0.5 μm. **c**, Experimental workflow showing the induction of *pGAL-HOP2/MND1* through the addition of β-estradiol at hour 4 of a meiotic time course after cells had formed heterologous SC. **d**, Immunostaining was performed with antibodies for Ecm11 and GFP at hours 4 (uninduced), 4.5, and 5 (after induction). DNA was counterstained with DAPI. **e**, Distance between GFP-dots in wild type at hour 4, induced *pGAL-HOP2/MND1* at hour 4, 4.5 and 5, and uninduced *pGAL-HOP2/MND1* at hour 4.5 and 5 as a measure of pairing. Box-and-whisker plots show medians and interquartile ranges. Grey dashed line indicates ≤ 0.5 μm, the cutoff used to consider dots paired. **f,** Quantification of percent of GFP-dots associated with Ecm11 signal in uninduced *pGAL-HOP2/MND1* at hour 4 (92.5 ± 6.4) and induced *pGAL-HOP2/MND1* at hour 4.5 (100 ± 0) and 5 (86.4 ± 3.46). **g**, Quantification of percent sporulation in wild type of monads (0.5%), dyads (10.5%) and tetrads (86.8%), *pGAL-HOP2/MND1* uninduced monads (2%), dyads (3%) and tetrads (0%) and induced monads (1%), dyads (14.5%) and tetrads (69.8%). **h**, Quantification of percent viable spores in wild type (97.5%), and induced *pGAL-HOP2/MND1* (98.38 ± 1.08%). The data presented are from chromosome spreads of synchronous meiotic yeast cultures and presented as mean ± s.d. Scale bars, 1 μm. Significance was evaluated using Mann-Whitney U test. n.s., not significant. **P ≤ 0.01.

To test whether heterologous synapsis can be corrected, we established an inducible system that delays strand invasion until heterologous synapsis is established. Initial attempts aimed at co-inducing Dmc1 and Rad51 failed to support gamete formation, likely because of the uneven stoichiometry of Dmc1 and Rad51 within the nucleoprotein filament, an interpretation supported by titration experiments (**Extended Data, Fig. 4**). Therefore, we instead focused on Hop2 because heterologous synapsis is well-established for *hop2Δ* mutants ^17^, and because Hop2 forms a stable dimer with Mnd1^44^, permitting balanced co-induction. We placed *HOP2* and *MND1* under the control of the *GAL* promoter and allowed cells to enter meiosis and undergo heterologous synapsis in the absence of Hop2/Mnd1 until hour 4, by which time maximal synapsis is established. We then induced expression of both proteins using the Gal4-ER expression system, which induces expression upon addition of β-estradiol ^45^, and assessed pairing of GFP-dots and synapsis over time using chromosome spreads (**Fig. 4c-e**). Prior to induction (T = 4 h), only 19% of the *pGAL-HOP2/MND1* nuclei showed pairing of GFP-dots compared to 92% of wild-type nuclei (**Fig. 4e**). This was unchanged at hour 4.5 following induction as uninduced and induced *pGAL-HOP2/MND1* cells displayed only 17% and 37% pairing respectively. However, by hour 5, levels of pairing increased to 76% in the *pGAL-HOP2/MND1* induced strain whereas the uninduced strain showed no major increase (**Fig. 4e**). Regardless of pairing status, most dots were associated with linear Ecm11 assemblies, implying synaptic engagement both prior and after induction (**Fig. 4f**). These data indicate that strand invasion can correct preexisting heterologous synapsis.

To test whether the corrected synapsis supports the formation of viable gametes, spore formation and gamete viability were assessed. Whereas uninduced *pGAL-HOP2/MND1* strains failed to form spores, tetrad formation in the induced strain reached close to wild-type levels (induced: 69.75% vs WT: 86.83%) (**Fig. 4g**). Moreover, spore viability was indistinguishable from wild type (induced: 98.4% vs WT: 97.4%) (**Fig. 4h**). Thus, initially heterologous synapsis did not prevent homologous chromosomes from pairing and segregating properly and hence is compatible with gamete formation.

Our results support the notion that chromosome synapsis is a dynamic process that can accommodate partner switches in response to meiotic recombination. Thus, although heterologous synapsis is a common phenotype of repair-deficient karyotypes, it is not a terminal pathology. The ability of chromosomal segments to change interaction partners is in line with the observation that individual SCs in *zip3* mutants can transiently disassembly ^19^ and the fact that synaptic adjustment allows chromosomal inversions to ultimately synapse with antiparallel, and thus non-homologous, chromosomal segments ^2,21^. Our results suggest that such alterations in synaptic partner choice can also occur on a much larger (i.e. nucleus-wide) scale to rectify incorrect synaptic partnerships. Although we did not observe overt nucleus-wide desynapsis during synaptic correction, desynapsis may be short-lived or occur asynchronously for different chromosomal segments. Alternatively, corrections may involve a more fluid fission and fusion of phase-separated structures within the SC.

We propose that heterologous synapsis reflects the natural ability of SCs to connect aligned chromosome axes independently of homology. During normal meiotic recombination, these initial connections are quickly corrected as homologous repair interactions identify matching sequences. The observation that heterologous synapsis in repair-deficient mutants exhibits characteristics of early stage SCs further implies that the identification of correct interactions promotes further SC maturation. Recombination-associated changes in SC dynamics have also been noted in *C. elegans* ^46^. SC assembly triggered by the crossover-associated synapsis initiation machinery (Zip2 and interactors) is an obvious candidate for this maturation step. It is possible that *MEC1*-dependent activation of synapsis provides an initial licensing step that restricts SC assembly until DSB processing has occurred, although the need for such licensing is debatable, as SC formation is uncoupled from DSB formation in several organisms ^2^. Alternatively, *MEC1*-dependent synapsis initiation may also have additional functions, as implied by our finding that the phosphomimetic mutant of Zip1-S75, which rescues synapsis, also improves spore viability in the *mec1-1* background. The nature of any such functions remains to be determined but our data identify SC formation as a dynamic process that interfaces at multiple points with meiotic recombination to ensure proper pairing and meiotic chromosome inheritance.

## Methods

### Yeast genetics, growth, and meiotic time-course assays

*S. cerevisiae* strains were generated with PCR-based methods ^47^ or CRISPR-based mutagenesis ^48^. All strains were of the SK1 background with the exception of the BR1919-8B strains shown in **Extended Data, Fig. 2e**. The strains used in this study are listed in **Extended Data, Table 1**. For synchronous meiosis of SK1 strains, cells were grown for approximately 24 hours in YPD at 25 °C, then diluted into BYTA (1% yeast extract, 2% bactotryptone, 1% potassium acetate, 50 mM potassium phthalate) at an OD_600_ of 0.3, grown overnight at 30 °C, then washed twice with water and resuspended in SPO medium (0.3% potassium acetate pH 7.0) at an OD_600_ of 1.9 at 30 °C to induce sporulation. Samples were taken at hours 0, 2, 3, 4, 5, 6 and processed. 5 mM β-estradiol were added to synchronous meiotic cultures at hour 4 in order to induce the expression of *HOP2* and *MND1* under the control of the *pGAL* promoter with strains expressing an Gal4-estradiol receptor fusion construct ^49^. Synchrony of all time courses was verified by flow cytometry (example shown in **Fig. 1f**); cells were fixed in ethanol, treated with proteinase K and RNAse prior to staining DNA with SYTOX Green and analyzing DNA content using a BD FACSAria in the Genomics Core at the NYU Center for Genomics and Systems Biology. For sporulation of BR strains, cells were grown in YPD until cultures reached an OD_600_ of 4-5. Cells from 1 volume of culture were then diluted in 5x volume of 2% potassium acetate to induce sporulation.

### Chromosome spreads

Meiotic cells were collected at hour 2, 3, 4, 4.5, 5, and 6 after meiotic induction and treated with 25 mM Tris/0.8M KCl pH 7.5/20 mM DTT for 2 min at room temperature and then spheroplasted in 25 mM Tris/0.8M KCl pH 7.5/ 0.13 μg/μL zymolyase T100 at 30 °C for 30 min. The spheroplasts were washed and resuspended in 0.1 M MES pH 6.4/1 mM EDTA/0.5 mM MgCl_2_/2 M Sorbitol. Cells were placed on a clean glass slide (soaked in ethanol and air-dried) and two volumes of fixative (3% paraformaldehyde/3.4% sucrose/2mM NaOH) were added to the cells and distributed across the slide by gentle tilting. Four volumes of 1% Lipsol (Bibby Sterlin Ltd.) were added to the slide and mixed by tilting. Four more volumes of fixative were added to the slide and then cells were spread by a clean glass rod. Slides were left to dry overnight and stored at −80 °C the next day. Prior to staining, slides were washed in 0.4% Kodak Photoflo (Cat no. 1464510), and blocked with 1× TBS/0.5% BSA/0.2% gelatin overnight at 4 °C. Slides were hybridized with antibodies at 37 °C for 1 hour (antibodies and corresponding dilutions are listed in **Extended Data, Table 2**). Slides were then washed with 1× TBS three times for 10 min each. Secondary antibodies were hybridized at 37 °C for 1 hour (**Extended Data, Table 2**). Slides were washed twice with 1× TBS and once with 0.4% Kodak Photoflo and mounted with DAPI mounting medium (Vectashield #H-1200-10). Stained slides were stored at 4 °C prior to imaging. Nuclei from meiotic BR cells were surface spread on glass slides and imaged as described in ^50^.

### Microscopy and cytological analysis

Images were collected on a Deltavision Elite imaging system (GE) equipped with an Olympus 100×/1.40 NA UPLSAPO PSF oil immersion lens and an InsightSSI Solid State Illumination module. Images were captured using an Evolve 512 EMCCD camera in the conventional mode and analyzed using ImageJ software. Scatterplots were generated using the GGplot2 package in R. STED images in **Fig. 1c** were taken using a Leica Stellaris confocal microscope with TauSTED mode with a 100x oil immersion lens at the Genomics Core at the New York University Center for Genomics and Systems Biology. The images in **Extended Data, Fig. 2e** were captured on a Leica SP8 STED microscope at The Stowers Institute. Images were deconvolved. The extent of synapsis was assessed by determining the total area of DAPI-associated SC protein signal or by measuring SC lengths, yielding identical results. By considering only DNA-associated signal, polycomplexes, which are aggregates of SC components that frequently form in DNA repair mutants and are not bound to DNA, could largely be excluded from quantification.

### Western blot analysis

For protein extraction, 5 mL of synchronous meiotic culture were harvested at hour 3 after meiotic induction. Cells were collected and resuspended in 5 mL of 5% trichloroacetic acid and incubated on ice for at least 10 min. Cell pellets were collected by centrifugation, washed with acetone and air dried overnight. Cells were ruptured with glass beads in 1× TE/250 mM DTT in a FastPrep-24 homogenizer. After adding SDS loading dye and adjusting to near neutral pH using Tris base, samples were boiled for 4 minutes, and then stored at -20°C. Proteins were separated by electrophoresis using a 7% polyacrylamide gel (150:1 acrylamide:bisacrylamide) in a Tris-acetate buffer. Proteins were blotted onto a nitrocellulose membrane (BioRad). The membrane was subsequently blocked with 5% milk in 1× PBS/0.1% Tween and incubated with rabbit anti-Zip1 antibody (y-300; Santa Cruz Biotechnology) at 1:1000 dilution overnight at 4°C. The membrane was washed three times with 1× PBS/0.1% Tween before incubation with anti-rabbit HRP secondary antibody (GE Healthcare Bio) for 2 hours at 4°C. The membrane was developed in Western Lightning-ECL substrate and imaged on Kodak film.

## Acknowledgements

We thank H. Klein and S. Keeney for helpful suggestions. We are indebted to S. Harrison for sharing the antibody against Ctf19 and G.S. Roeder for the antibody against Red1. We thank the Stowers Institute for use of their Leica SP8 STED microscope. We acknowledge the Genomics Core at the New York University Center for Genomics and Systems Biology for access to instrumentation and valuable technical assistance, and are grateful to the Zegar Family Foundation for their generous support of the NYU Genomics Core.

## Author contributions

Conceptualization – L.M.L. and A.H.; Investigation & formal analysis – L.M.L., A.J.M., C.M., A.H.; Essential reagents – A.J.M., N.M.H.; Manuscript writing (initial draft) – L.M.L. and A.H.; Manuscript editing – all authors.

---

**Extended Data, Figure 1:**
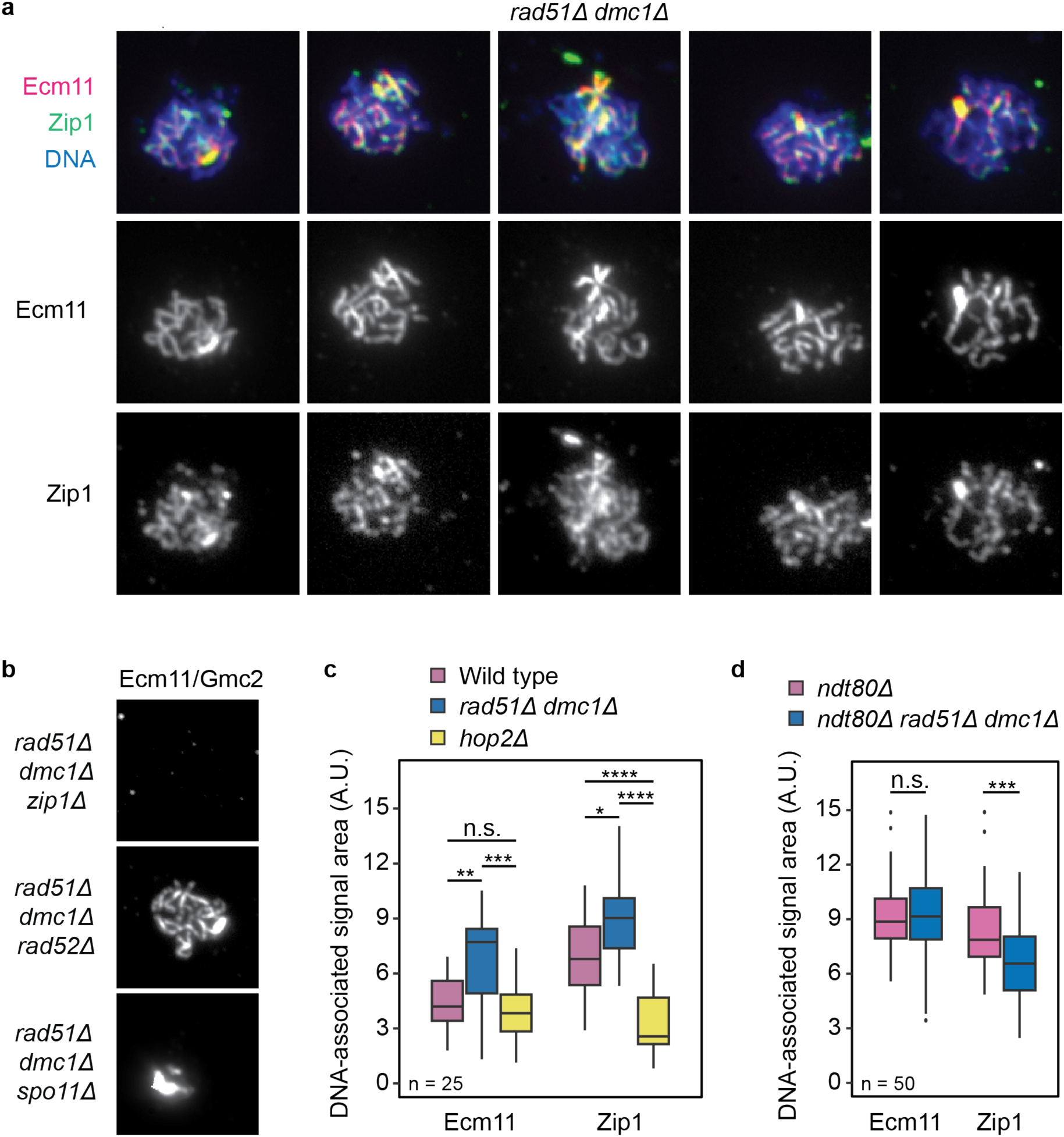
Characterization of synapsis in recombination mutants. **a**, Additional example spread nuclei from a synchronous culture of *rad51Δ dmc1Δ* showing extensive synapsis at hour 4. Immunostaining was performed with antibodies for Ecm11 and Zip1. DNA was counterstained with DAPI. **b**, Spread nuclei of *rad51Δ dmc1Δ* cells also lacking *ZIP1*, *RAD52*, or *SPO11*. Immunostaining was performed with antibodies for Ecm11 or Gmc2 at hour 4. **c**, Quantification of DNA-associated signal area (A.U.) per cell at hour 4 in wild type (Gmc2; 4.50 ± 1.48, Zip1; 6.76 ± 2.12), *rad51Δ dmc1Δ* (Gmc2; 6.68 ± 2.54, Zip1; 8.49 ± 3.26), and *hop2Δ* (Gmc2; 3.95 ± 1.57, Zip1; 3.60 ± 2.08). **d**, Quantification of DNA-associated signal area (A.U.) per cell at hour 6 in *ndt80Δ* (Ecm11; 9.09 ± 1.87, Zip1; 8.29 ± 2.18) and *ndt80Δ rad51Δ dmc1Δ* (Ecm11; 9.29 ± 2.53, Zip1; 6.67 ± 2.19). The data presented are from chromosome spreads of synchronous meiotic yeast cultures and presented as mean ± s.d. Scale bars, 1 μm. Significance was evaluated using Mann-Whitney U test. n.s., not significant. *P ≤ 0.05, **P ≤ 0.01, ***P ≤ 0.001, ****P ≤ 0.0001.

**Extended Data, Figure 2:**
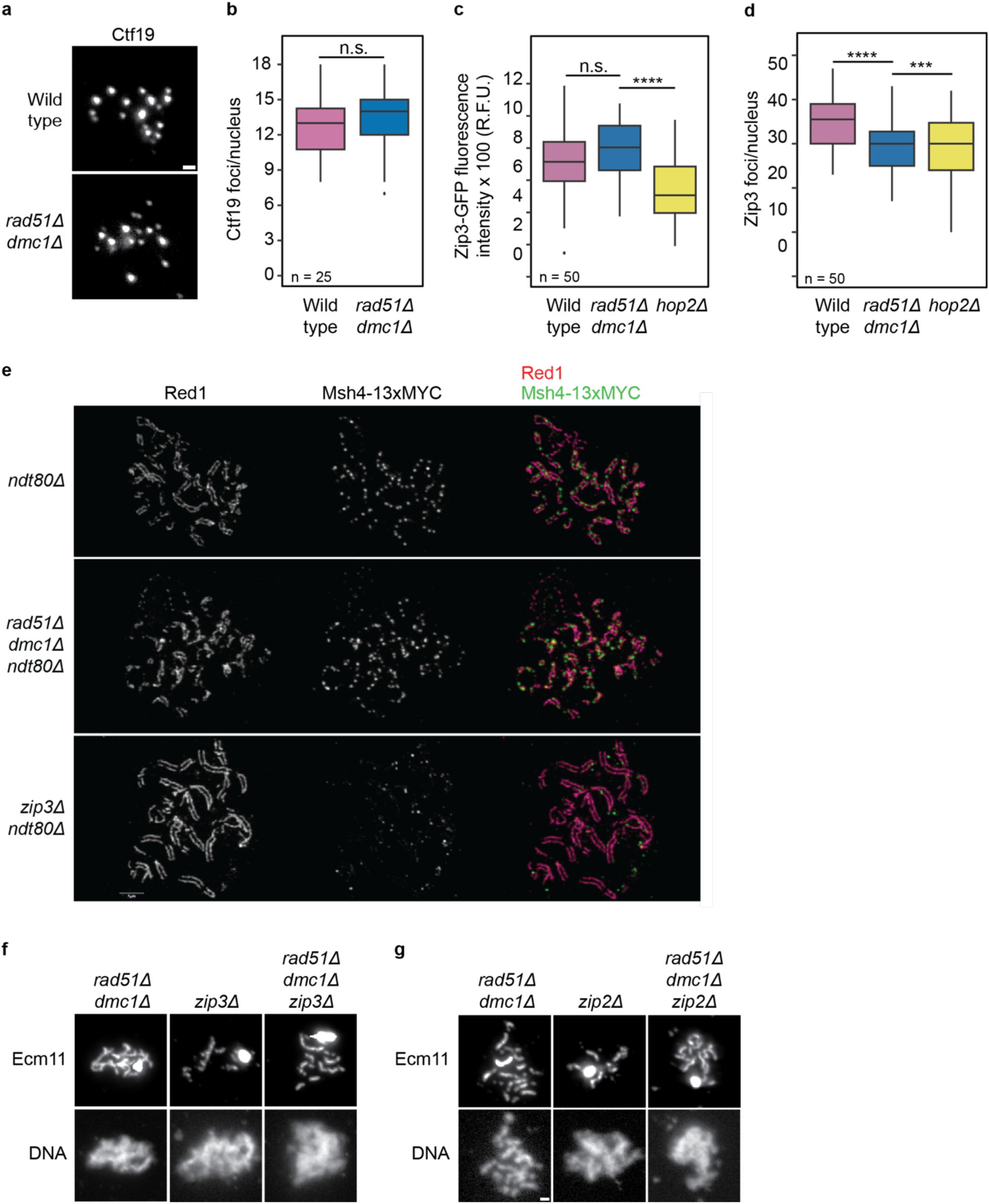
Centromeres pair and canonical markers of crossover designation are retained in rad51Δ dmc1Δ mutants. **a**, Spread nuclei of wild type and *rad51Δ dmc1Δ* cells. Immunostaining was performed with an antibody for Ctf19 at hour 3. **b**, Quantification of the number of Ctf19 foci per nucleus at hour 3 in wild type (12.70 ± 2.58) and *rad51Δ dmc1Δ* (13.40 ± 3.13). **c**, Quantification of Zip3-GFP fluorescence intensity x 100 (R.F.U.) at hour 4 in wild type (726 ± 264), *rad51Δ dmc1Δ* (793 ± 98), and *hop2Δ* (542 ± 203). **d**, Quantification of the number of Zip3 foci per nucleus at hour 4 in wild type (35.4 ± 9.71), *rad51Δ dmc1Δ* (29.1 ± 5.50), and *hop2Δ* (29.4 ± 6.97). **e**, Spread nuclei of *ndt80Δ*, *rad51Δ dmc1Δ ndt80Δ* and *zip3Δ ndt80Δ* cells carrying a Msh4-13xMYC construct. Immunostaining was performed with antibodies for Red1 and MYC at 24 hours. Samples were imaged using STED microscopy. **f**, Spread nuclei of meiotic *rad51Δ dmc1Δ, zip3Δ*, and *rad51Δ dmc1Δ zip3Δ* cells. Immunostaining was performed with an antibody for Ecm11 at hour 4. DNA was counterstained with DAPI. **g**, Spread nuclei of meiotic *rad51Δ dmc1Δ, zip2Δ*, and *rad51Δ dmc1Δ zip2Δ* cells. Immunostaining was performed with an antibody for Ecm11 at hour 4. DNA was counterstained with DAPI. The data presented are from chromosome spreads of synchronous meiotic yeast cultures and presented as mean ± s.d. Scale bars, 1 μm. Significance was evaluated using Mann-Whitney U test. n.s., not significant. ***P ≤ 0.001, ****P ≤ 0.0001.

**Extended Data, Figure 3:**
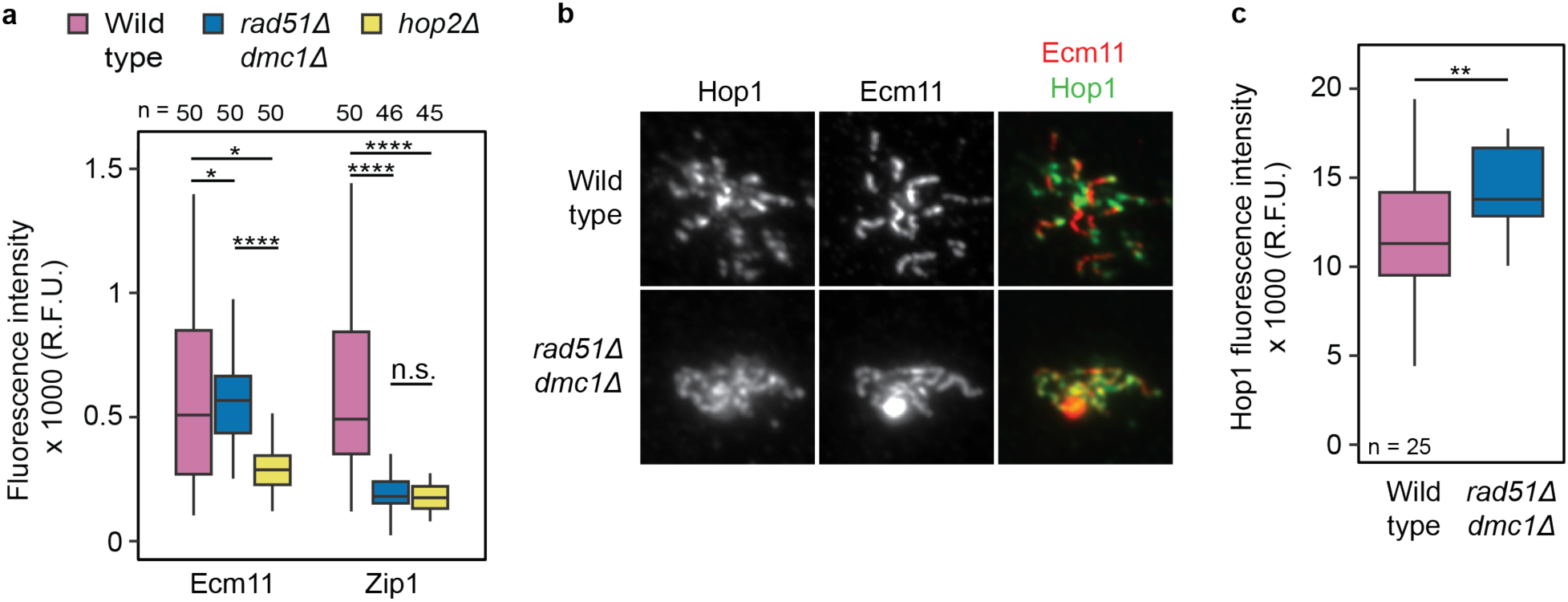
Synapsis in rad51Δ dmc1Δ mutant displays features of early prophase. **a**, Quantification of fluorescence intensity x 1000 (R.F.U.) (excluding polycomplexes) in wild type (Ecm11; 597 ± 375, Zip1; 626 ± 376), *rad51Δ dmc1Δ* (Ecm11; 585 ± 235, Zip1; 190 ± 66.10), and *hop2Δ* (Ecm11; 305 ± 180, Zip1; 169 ± 50.10). The data presented are from chromosome spreads of synchronous meiotic yeast cultures and presented as mean ± s.d. Scale bars, 1 μm. Significance was evaluated using Mann-Whitney U test. n.s., not significant. *P ≤ 0.05, **P ≤ 0.01, ***P ≤ 0.001, ****P ≤ 0.0001.**b**, Spread nuclei of meiotic wild type and *rad51Δ dmc1Δ* cells. Immunostaining was performed with antibodies for Ecm11 and Hop1 at hour 5. DNA was counterstained with DAPI. **c**, Quantification of Hop1 fluorescence intensity x 1000 (R.F.U.) at hour 5 in wild type (11.88 ± 3.91) and *rad51Δ dmc1Δ* (15.07 ± 3.26). The data presented are from chromosome spreads of synchronous meiotic yeast cultures and presented as mean ± s.d. Scale bars, 1 μm. Significance was evaluated using Mann-Whitney U test. n.s., not significant. **P ≤ 0.01, ***P ≤ 0.001, ****P ≤ 0.0001.

**Extended Data, Figure 4:**
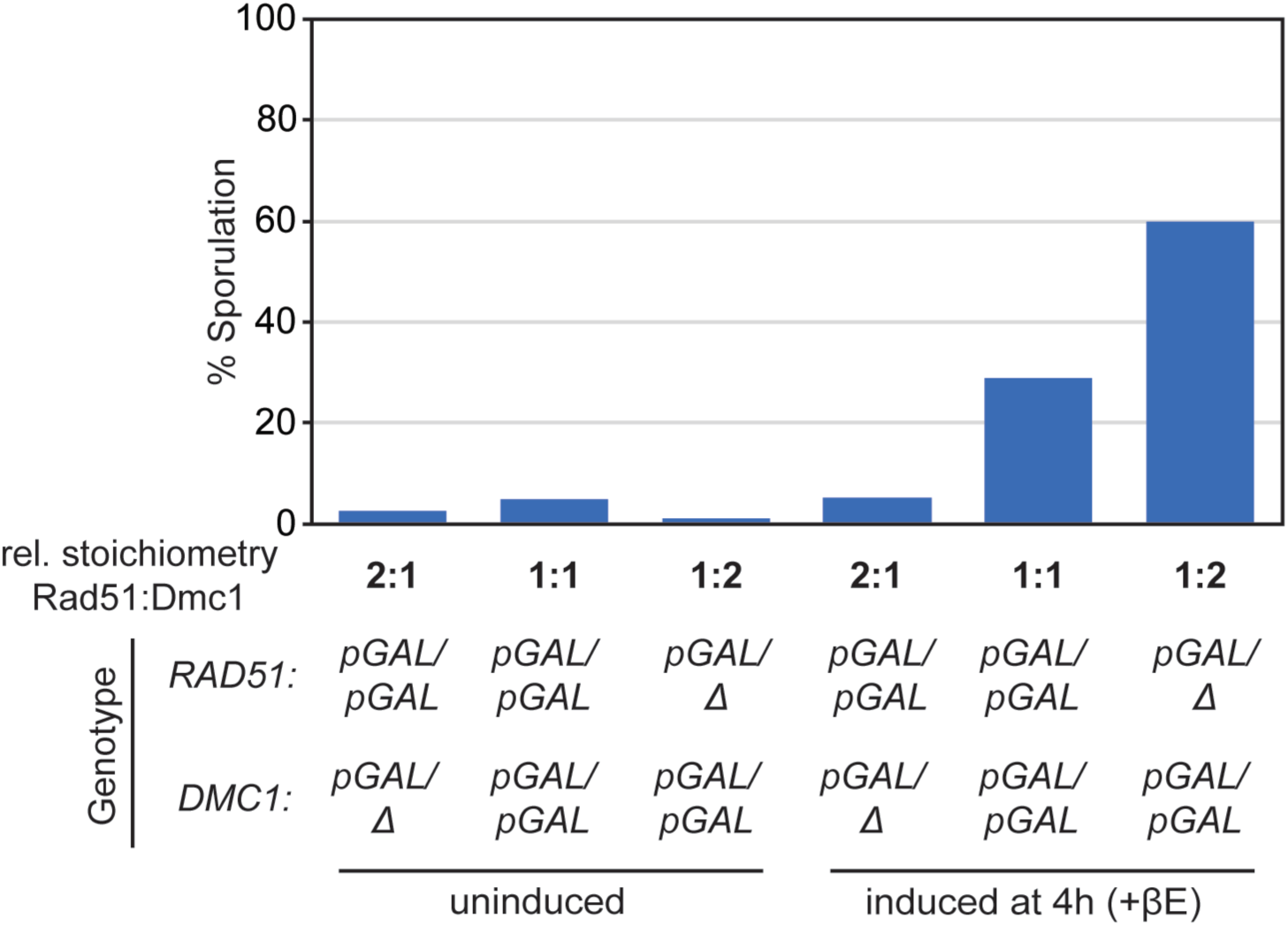
Sporulation is differentially affected by RAD51/DMC1 stoichiometry. Quantification of percent sporulation for *pGAL-RAD51* and/or *pGAL-DMC1* strains. Strains carried either 1 or 2 copies of the induction constructs, resulting in different relative stoichiometries of *RAD51* and *DMC1* expression. The relative Rad51:Dmc1 stoichiometry upon induction affected sporulation efficiency: uninduced 2:1 (2.5%), 1:1 (5%), 1:2 (1%) and induced at hour 4 with β-estradiol 2:1 (5%), 1:1 (29%), 1:2 (60%).

## Extended Data, Tables

**Extended Data, Table 1:**
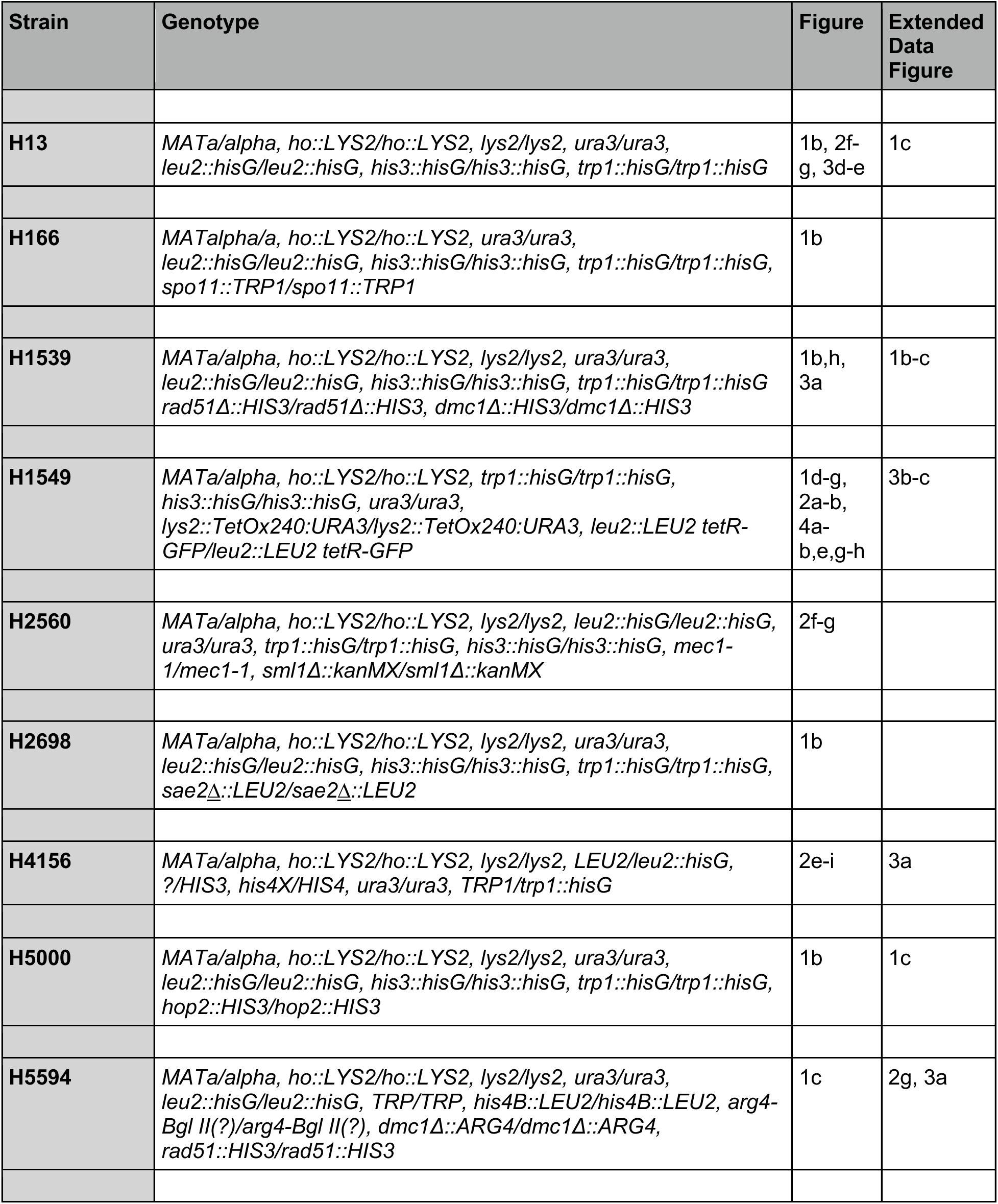
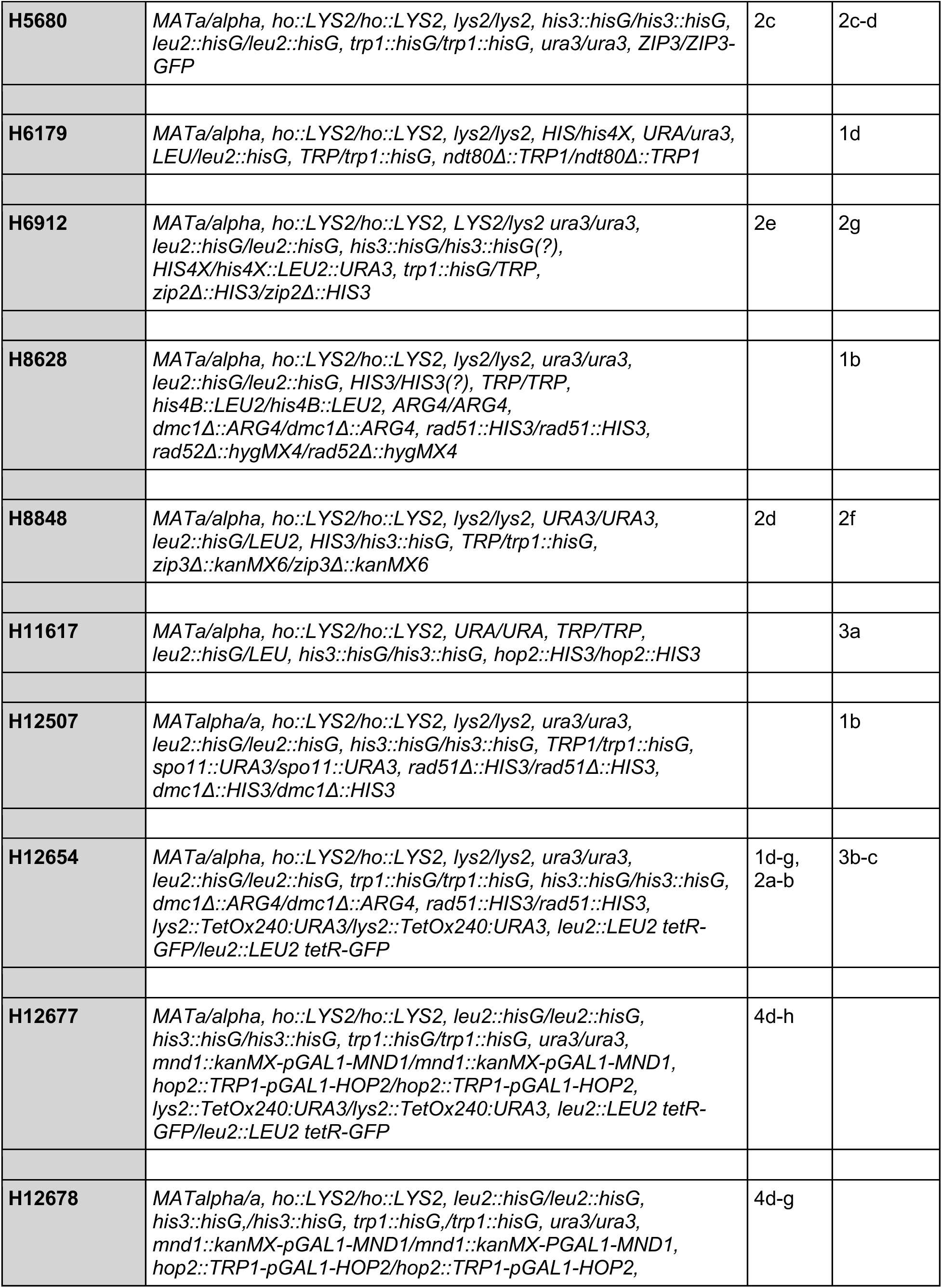
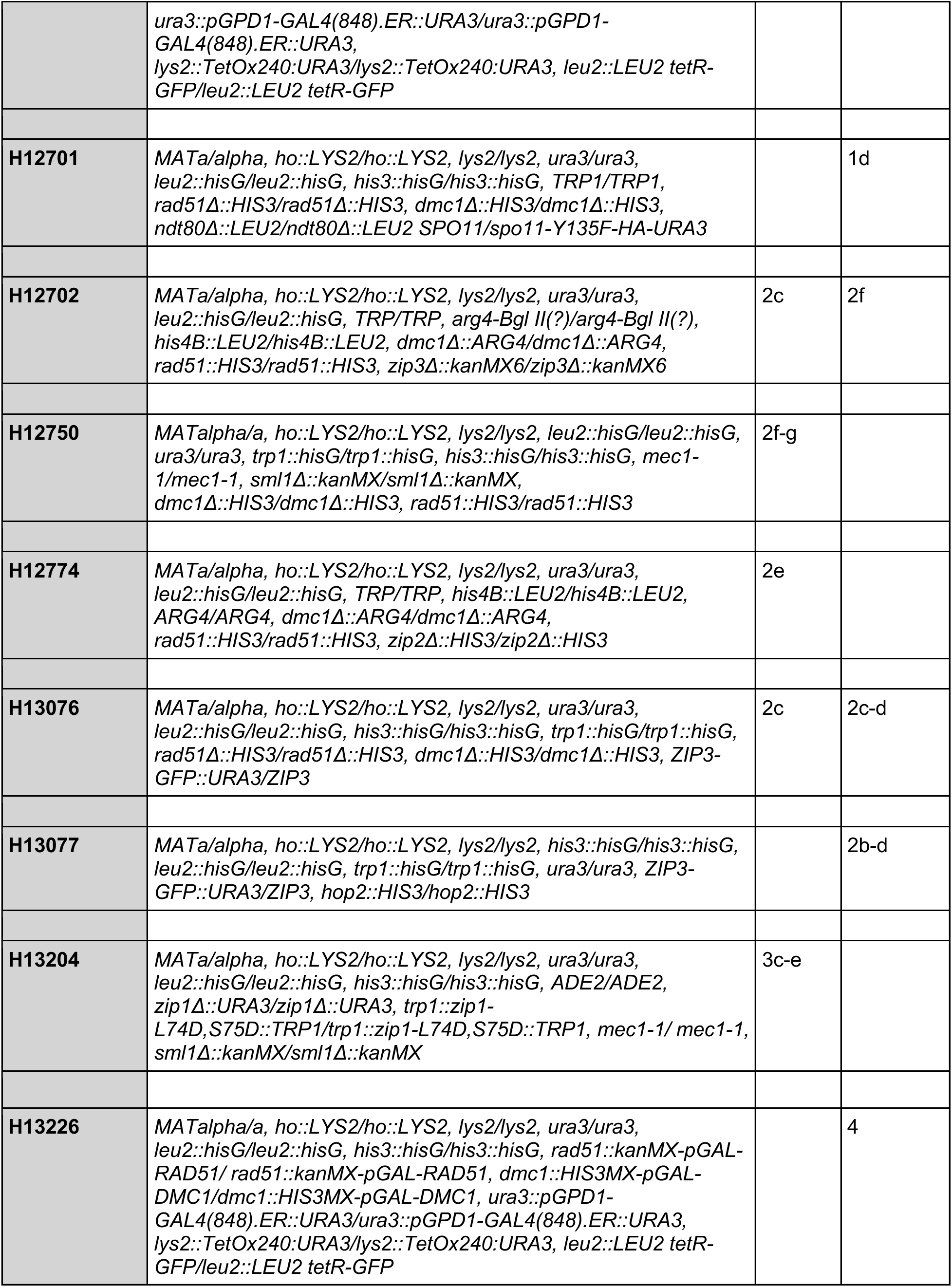
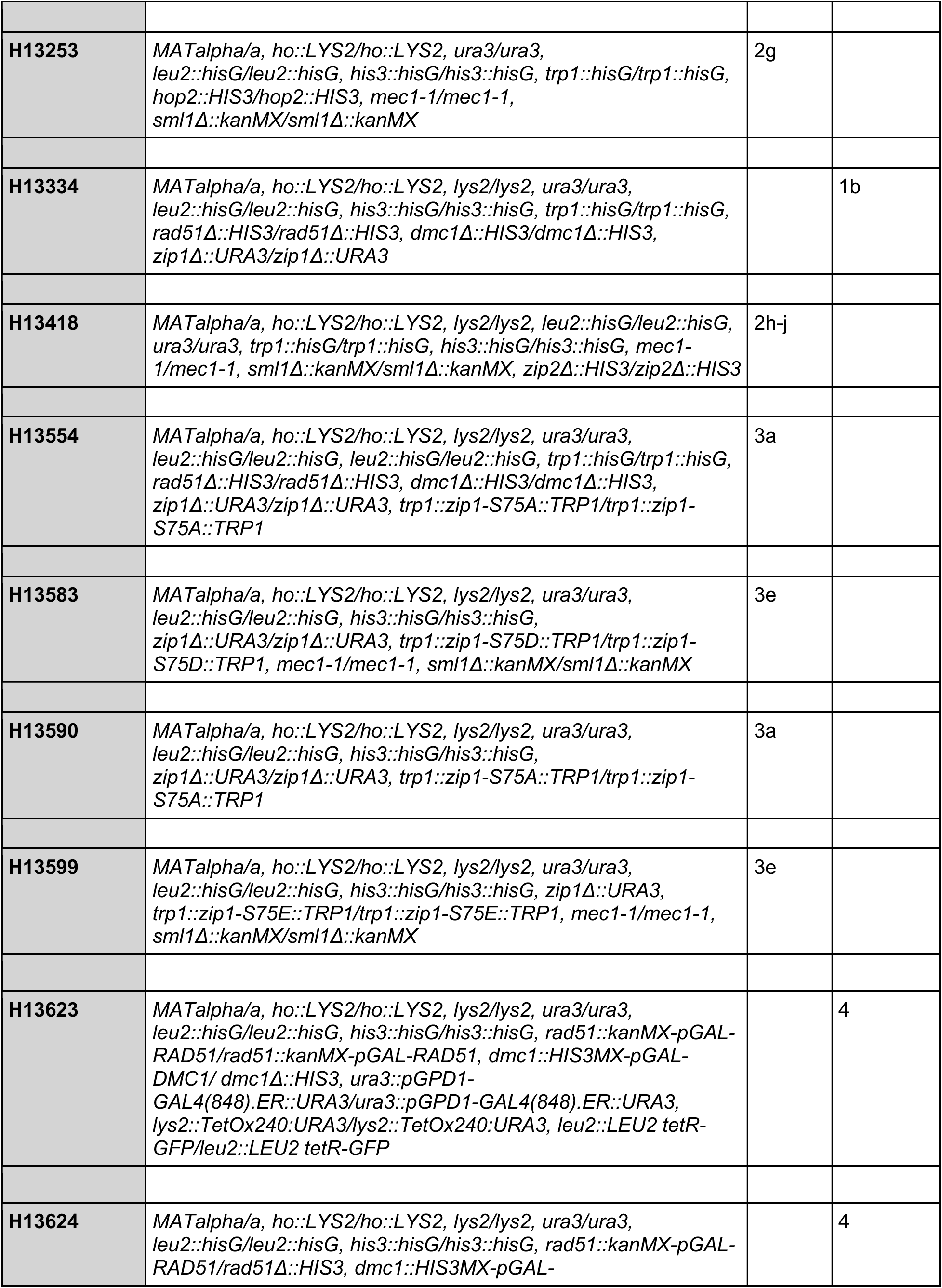
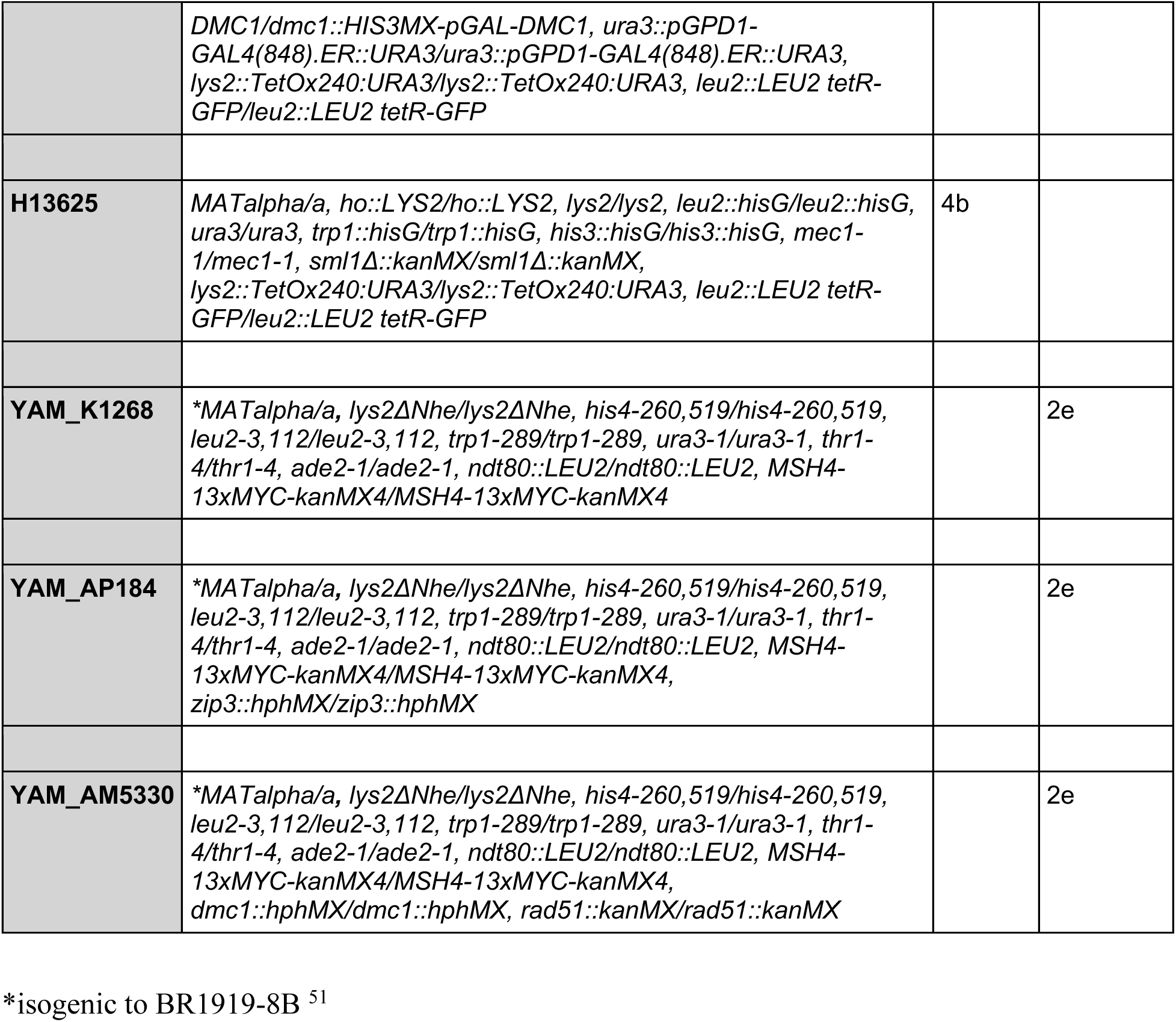
Strains used in this study.

**Extended Data, Table 2:**
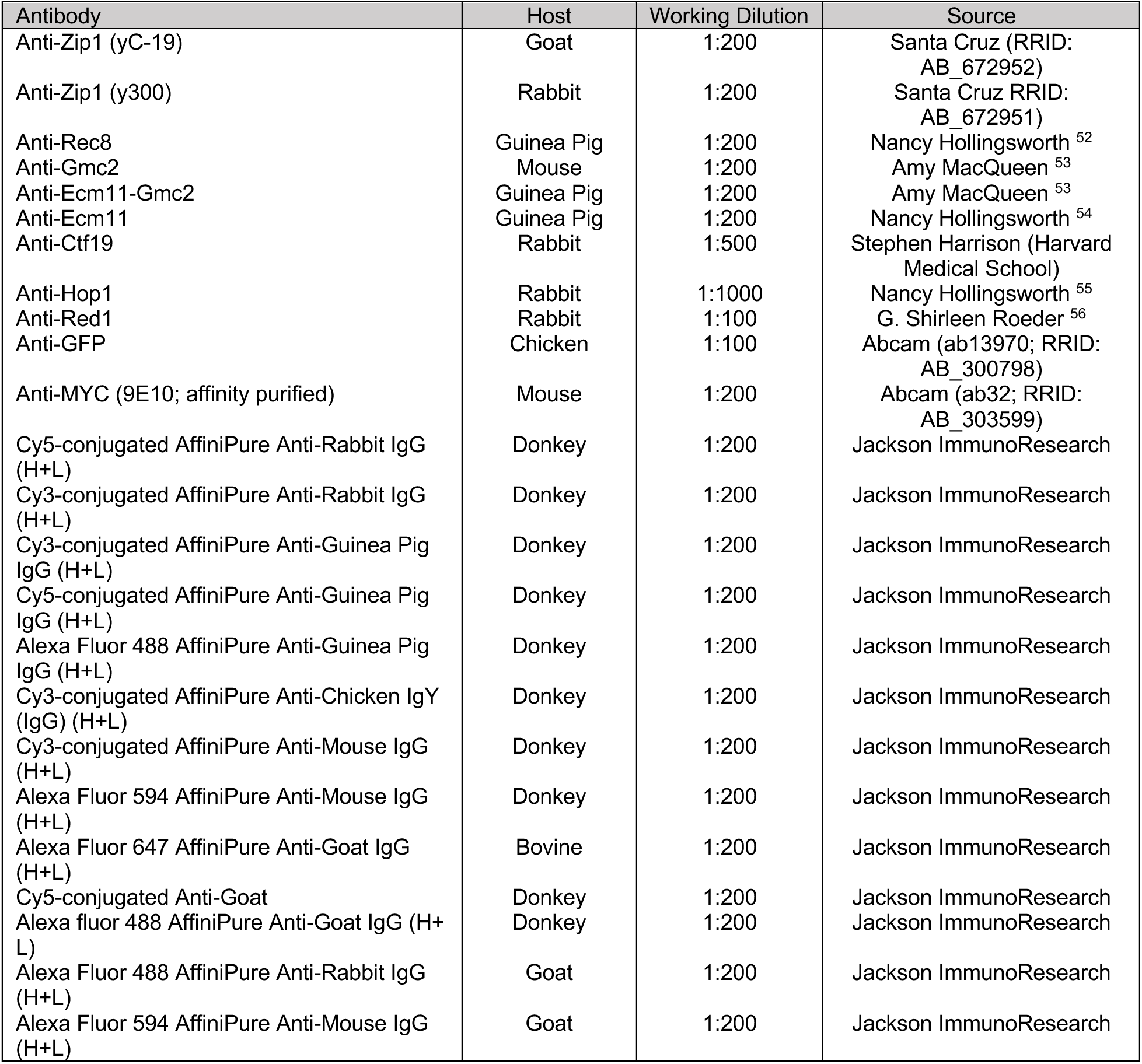
Immuno-staining conditions used in this study.

| Antibody | Host | Working Dilution | Source |
| --- | --- | --- | --- |
| Anti-Zip1 (yC-19) | Goat | 1:200 | Santa Cruz (RRID: AB_672952) |
| Anti-Zip1 (y300) | Rabbit | 1:200 | Santa Cruz RRID: AB_672951) |
| Anti-Rec8 | Guinea Pig | 1:200 | Nancy Hollingsworth <sup>52</sup> |
| Anti-Gmc2 | Mouse | 1:200 | Amy MacQueen <sup>53</sup> |
| Anti-Ecm11-Gmc2 | Guinea Pig | 1:200 | Amy MacQueen <sup>53</sup> |
| Anti-Ecm11 | Guinea Pig | 1:200 | Nancy Hollingsworth <sup>54</sup> |
| Anti-Ctf19 | Rabbit | 1:500 | Stephen Harrison (Harvard Medical School) |
| Anti-Hop1 | Rabbit | 1:1000 | Nancy Hollingsworth <sup>55</sup> |
| Anti-Red1 | Rabbit | 1:100 | G. Shirleen Roeder <sup>56</sup> |
| Anti-GFP | Chicken | 1:100 | Abcam (ab13970; RRID: AB_300798) |
| Anti-MYC (9E10; affinity purified) | Mouse | 1:200 | Abcam (ab32; RRID: AB_303599) |
| Cy5-conjugated AffiniPure Anti-Rabbit IgG (H+L) | Donkey | 1:200 | Jackson ImmunoResearch |
| Cy3-conjugated AffiniPure Anti-Rabbit IgG (H+L) | Donkey | 1:200 | Jackson ImmunoResearch |
| Cy3-conjugated AffiniPure Anti-Guinea Pig IgG (H+L) | Donkey | 1:200 | Jackson ImmunoResearch |
| Cy5-conjugated AffiniPure Anti-Guinea Pig IgG (H+L) | Donkey | 1:200 | Jackson ImmunoResearch |
| Alexa Fluor 488 AffiniPure Anti-Guinea Pig IgG (H+L) | Donkey | 1:200 | Jackson ImmunoResearch |
| Cy3-conjugated AffiniPure Anti-Chicken IgY (IgG) (H+L) | Donkey | 1:200 | Jackson ImmunoResearch |
| Cy3-conjugated AffiniPure Anti-Mouse IgG (H+L) | Donkey | 1:200 | Jackson ImmunoResearch |
| Alexa Fluor 594 AffiniPure Anti-Mouse IgG (H+L) | Donkey | 1:200 | Jackson ImmunoResearch |
| Alexa Fluor 647 AffiniPure Anti-Goat IgG (H+L) | Bovine | 1:200 | Jackson ImmunoResearch |
| Cy5-conjugated Anti-Goat | Donkey | 1:200 | Jackson ImmunoResearch |
| Alexa fluor 488 AffiniPure Anti-Goat IgG (H+L) | Donkey | 1:200 | Jackson ImmunoResearch |
| Alexa Fluor 488 AffiniPure Anti-Rabbit IgG (H+L) | Goat | 1:200 | Jackson ImmunoResearch |
| Alexa Fluor 594 AffiniPure Anti-Mouse IgG (H+L) | Goat | 1:200 | Jackson ImmunoResearch |

